# *In vivo* and *in vitro* analysis of the role of the Prc protease in inducing mucoidy in *Pseudomonas aeruginosa*

**DOI:** 10.1101/2024.05.28.596254

**Authors:** Alexis G. Sommerfield, Michelle Wang, Julia Mamana, Andrew J. Darwin

**Author notes:** Address correspondence to: Andrew J. Darwin. co-first author (Alexis G. Sommerfield, Michelle Wang).

## Abstract

In *Pseudomonas aeruginosa,* alginate biosynthesis gene expression is inhibited by the transmembrane anti-sigma factor MucA, which sequesters the AlgU sigma factor. Cell envelope stress initiates cleavage of the MucA periplasmic domain by site-1 protease AlgW, followed by further MucA degradation to release AlgU. However, after colonizing the lungs of people with cystic fibrosis, *P. aeruginosa* converts to a mucoid form that produces alginate constitutively. Mucoid isolates often have *mucA* mutations, with the most common being *mucA22*, which truncates the periplasmic domain. MucA22 is degraded constitutively, and genetic studies suggested that the Prc protease is responsible. Some studies also suggested that Prc contributes to induction in strains with wild type MucA, whereas others suggested the opposite. However, missing from all previous studies is a demonstration that Prc cleaves any protein directly, which leaves open the possibility that the effect of a *prc* null mutation is indirect. To address the ambiguities and shortfalls, we reevaluated the roles of AlgW and Prc as MucA and MucA22 site-1 proteases. *In vivo* analyses using three different assays, and two different inducing conditions, all suggested that AlgW is the only site-1 protease for wild type MucA in any condition. In contrast, genetics suggested that AlgW or Prc act as MucA22 site-1 proteases in inducing conditions, whereas Prc is the only MucA22 site-1 protease in non-inducing conditions. For the first time, we also show that Prc is unable to degrade the periplasmic domain of wild type MucA, but does degrade the mutated periplasmic domain of MucA22 directly.

**IMPORTANCE:** After colonizing the lungs of individuals with cystic fibrosis, *P. aeruginosa* undergoes mutagenic conversion to a mucoid form, worsening the prognosis. Most mucoid isolates have a truncated negative regulatory protein MucA, which leads to constitutive production of the extracellular polysaccharide alginate. The protease Prc has been implicated, but not shown, to degrade the most common MucA variant, MucA22, to trigger alginate production. This work provides the first demonstration that the molecular mechanism of Prc involvement is direct degradation of the MucA22 periplasmic domain, and perhaps other truncated MucA variants as well. MucA truncation and degradation by Prc might be the predominant mechanism of mucoid conversion in cystic fibrosis infections, suggesting that Prc activity could be a useful therapeutic target.

## INTRODUCTION

The Gram-negative bacterium *Pseudomonas aeruginosa* is a significant health care burden, especially as a common cause of antibiotic-resistant infections in hospitals, and in those suffering from Cystic fibrosis (CF) (1). CF is caused by a defective CFTR chloride ion transporter that causes multiple symptoms, including mucus buildup in the respiratory system (2, 3). CF lungs often become colonized by *P. aeruginosa*, which has been the leading cause of morbidity and mortality (4). After colonizing the lungs, *P. aeruginosa* converts to a form that overproduces the extracellular polysaccharide alginate constitutively (5). These mucoid *P. aeruginosa* strains are difficult to eradicate, in part because the thick alginate coat protects them from antibiotics and components of the host immune response (6–8).

Expression of the *algD* operon, which encodes most of the proteins required for alginate production and export, is controlled by the extracytoplasmic function sigma factor AlgU (also known as AlgT) (9–12). AlgU is 66% similar to *E. coli* σ^E^ (RpoE), which controls a cell envelope stress response (12–15). σ^E^ is negatively regulated by RseA, a transmembrane anti-sigma factor, with a cytoplasmic N-terminal domain that sequesters σ^E^ and a C-terminal periplasmic domain. Another protein named RseB binds to the periplasmic domain of RseA and protects it from the site-1 protease DegS. Two signals initiate regulated intramembrane proteolysis of RseA: mislocalized LPS molecules interact with RseB to dissociate it from the RseA periplasmic domain, and mislocalized outer membrane proteins (OMPs) interact with DegS to induce it to cleave the unprotected periplasmic domain of RseA (16). After DegS cleavage, the truncated RseA is a substrate for the intramembrane protease RseP/YaeL, which cleaves the RseA transmembrane domain. The residual cytoplasmic region of RseA is degraded by the ClpXP protease, which frees σ^E^ to bind to target promoters (15, 17–20).

*P. aeruginosa* AlgU, MucA and MucB are functionally analogous to the *E. coli* σ^E^, RseA and RseB proteins, respectively (21). MucB protects the C-terminus of MucA from cleavage by the site-1 protease, named AlgW in *P. aeruginosa*. MucB is released from MucA by mislocalized LPS, and the AlgW protease is activated by mislocalized OMPs (16, 22, 23). AlgW cleaves MucA between Ala-136 and Gly-137 (23). MucA is then further degraded similarly to RseA in *E. coli*, releasing the AlgU sigma factor to induce target genes, including those that promote the production and export of alginate.

Constitutively mucoid *P. aeruginosa* strains from the majority of CF patients have *mucA* mutations (24–30). Many of these cause a C-terminal truncation of MucA, and one of the most common is the *mucA22* allele (24–27, 31). *mucA22* is a G-C base pair deletion, which causes a frameshift after codon Gly-143 that adds three arginine residues followed by a premature stop codon (24). The mucoid phenotype is thought to be caused by constitutive degradation of MucA22 to release AlgU. Suppressor screens revealed that null mutations in *prc* (also called *algO*) reverted *mucA22* strains to a nonmucoid phenotype (32, 33). This suggested that Prc, which is a carboxyl-terminal processing protease, might be responsible for constitutive MucA22 degradation (34). However, direct evidence for this has not been reported.

The original Prc-MucA study indicated that Prc promoted mucoidy in *mucA22* strains, but might not act against wild type MucA (32). In agreement with that, the induction of a mucoid phenotype by irgasan and ammonium metavanadate in a wild type MucA strain was also unaffected by Prc (35). However, a later report found that Prc was required to induce a mucoid phenotype in those conditions (36). A *prc* null mutation was also reported to reduce D-cycloserine-dependent induction of the AlgU regulon in a *mucA*^+^ strain, although it was later determined that the effect was minor (37, 38). Finally, one group reported that a *prc* null mutation abolished induction of the AlgU regulon in response to MucE overproduction in a *mucA*^+^ strain, whereas another group reported earlier that Prc was not required for MucE-dependent induction (39, 40). The ambiguity described above leaves open the possibility that Prc could cleave both MucA22 and wild type MucA. Indeed, Prc has been proposed to cleave some other wild type transmembrane anti-sigma factors in *Pseudomonas* species, including *P. aeruginosa* (41–44). However, missing from all of those studies is experimental evidence that Prc cleaves any protein directly, including MucA22. In *E. coli,* Prc degrades peptidoglycan lytic enzymes, and in *P. aeruginosa* cell wall stress is known to affect the AlgU regulon (37, 45–49). Therefore, altered cell wall metabolism in a *prc* null mutant could interfere with activation of the AlgU regulon indirectly, in MucA and/or MucA22 strains. Here, we report a focused set of systematic experiments to address all of these issues with an assessment and comparison of the roles of AlgW and Prc as site-1 proteases for MucA and MucA22. We used three different assays, and two different inducing conditions, to monitor activation of the AlgU regulon qualitatively and quantitatively *in vivo*. We also purified proteins to test the ability of Prc to cleave the C-terminal domains of MucA and MucA22 directly. For the first time, our data demonstrate conclusively that Prc can act as a constitutive site-1 protease for MucA22, but it cannot degrade the periplasmic domain of the wild type MucA protein.

## RESULTS

### The effects of two carboxyl-terminal processing proteases on the induction of alginate biosynthesis

Prc is a member of the carboxyl-terminal processing protease (CTP) family. There is one other CTP in *P. aeruginosa*, named CtpA (34). There has not been any investigation into the effect of CtpA on the AlgU regulon. Therefore, we began by comparing the effects of both CTPs on AlgU regulon activation, in *mucA^+^* and *mucA22* strains. We used Δ*prc* and Δ*ctpA* in-frame deletion mutations, two different inducing conditions, and three different phenotypic readouts for AlgU activation.

The first condition we used was the peptidoglycan synthesis inhibitor D-cycloserine, which has been shown to induce the AlgU regulon in a MucA-degradation dependent manner (37). We monitored Φ(*algDp-lacZ*) operon fusion expression by quantitative β-Galactosidase assay, and endogenous AlgD protein levels by qualitative immunoblot. Without D-cycloserine, there was low Φ(*algDp-lacZ*) expression and undetectable levels of AlgD protein in all *mucA*^+^ strains (wild type, Δ*prc*, Δ*ctpA*; Fig. 1a,b). Exposure of the wild type strain to D-cycloserine increased Φ(*algDp-lacZ*) expression approximately 100-fold, and the AlgD protein was now detected (Fig. 1a,b). Neither a Δ*prc* or Δ*ctpA* mutation led to reduced Φ(*algDp-lacZ*) expression or AlgD abundance. Therefore, these data confirm that CtpA and Prc are both not required for induction of the AlgU regulon when wild type MucA is present. We did notice that a Δ*ctpA* mutation slightly increased both Φ(*algDp-lacZ*) expression and AlgD protein level in the presence of D-cycloserine (Fig. 1a,b). This might be due to Δ*ctpA* exacerbating cell wall stress (see Discussion).

**FIG 1.**
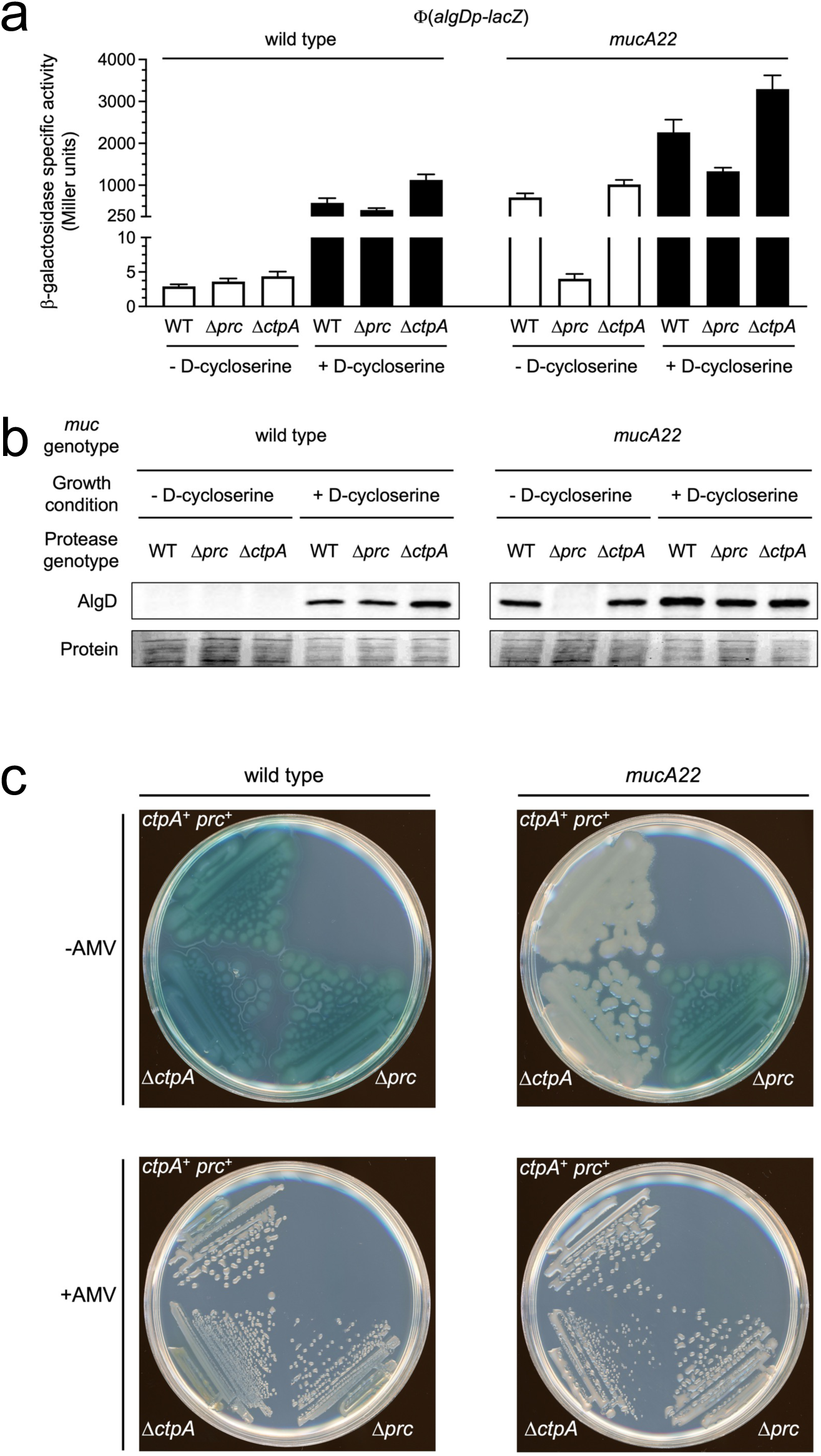
Effects of Δ*prc* and Δ*ctpA* mutations on regulation of the *algD* operon and mucoidy. (a) Φ(*algDp-lacZ*) operon fusion expression. Strains with the wild type *mucA* gene (wild type) or *mucA22* were grown with or without exposure to D-cycloserine. Protease genotypes are shown below each bar (WT = wild type *prc*^+^ *ctpA*^+^). Error bars indicate the positive standard deviations from the means. (b) Anti-AlgD immunoblot analysis of one representative set of the triplicate cultures used to generate the data in panel a. (c) Phenotypes on Pseudomonas Isolation Agar with or without ammonium metavanadate (AMV). Immunoblots and agar plates are single representatives of several replicate experiments.

In the absence of D-cycloserine, a *mucA22* mutation increased the Φ(*algDp-lacZ*) expression and AlgD protein levels to a similar extent as induction with D-cycloserine had done in the *mucA*^+^ strain (Fig. 1a,b). Both phenotypes were completely suppressed by Δ*prc*, but not by Δ*ctpA* (Fig. 1a,b). This confirmed previous reports that Prc is required for the constitutive *mucA22* phenotype. Exposure to D-cycloserine restored Φ(*algDp-lacZ*) expression and AlgD protein production in the *mucA22* Δ*prc* strain, showing that Prc is not required for signal-responsive induction with MucA22. The Δ*ctpA* mutation slightly increased Φ(*algDp-lacZ*) expression in the presence of D-cycloserine in a *mucA22* strain, as it had done in the *mucA*^+^ strain (Fig. 1a,b).

Next, we used a different inducing condition and phenotypes to corroborate our findings. Growth on *Pseudomonas* isolation agar (PIA) supplemented with the phosphatase inhibitor ammonium metavanadate (AMV) induces a mucoid phenotype (35). The physiological relevance of these inducing conditions is unclear, but they do induce the normal signal transduction pathway that results in MucA degradation (35). When *mucA^+^* strains were grown on PIA without AMV, they appeared blue/green, presumably due to pyocyanin production, and non-mucoid, regardless of their *prc* and *ctpA* genotype (Fig. 1c). When AMV was added, the *mucA*^+^ strains became mucoid, as evidenced by the shiny/moist appearance of the colonies (Fig. 1c). They also lost the blue/green pigment, consistent with a report that activation of the AlgU regulon coincides with reduced pyocyanin production prior to stationary phase (31). These observations confirm the findings with D-cycloserine induction, showing that neither Prc or CtpA is required for induction of the AlgU regulon and mucoidy with wild type MucA. As expected, the *mucA22* mutation caused a mucoid phenotype in the absence of AMV, which was suppressed by Δ*prc* but not by Δ*ctpA* (Fig. 1c). In the presence of AMV, all three *mucA22* strains were mucoid (Fig. 1c). Again, these findings supported the conclusions made from the D-cycloserine experiments (Fig. 1a,b).

These experiments show that neither of the *P. aeruginosa* CTPs plays a significant role in activating the AlgU regulon and mucoidy in *mucA^+^* strains. They also validated previous work showing that Prc is required for the constitutive induction in a *mucA22* strain. However, Prc becomes dispensable for MucA22-dependent induction in the presence of inducing signals, presumably because AlgW is now activated and can take on the role as the MucA22 site-1 protease. Therefore, next we compared the contributions of the two proposed site-1 proteases, AlgW and Prc.

### *In vivo* analysis of AlgW- and Prc-dependent activation of the AlgU regulon in strains with MucA or MucA22

We compared the phenotypes of wild type MucA strains with *Δprc*, *ΔalgW* or *Δprc ΔalgW* in-frame deletion mutations. There was minimal Φ(*algDp-lacZ*) expression in the absence of D-cycloserine in the *mucA*^+^ strains, and the AlgD protein was not detected (Fig. 2a,b). Exposure to D-cycloserine increased Φ(*algDp-lacZ*) expression and AlgD protein production in wild type and Δ*prc* strains, as observed in the previous experiments (compare Fig. 2a,b to Fig. 1a,b). However, this induction was essentially abolished in *ΔalgW* and *Δprc ΔalgW* mutants, which is consistent with AlgW being the site-1 protease for wild type MucA (Fig. 2a,b). There was a small residual D-cycloserine-dependent increase in Φ(*algDp-lacZ*) expression in the Δ*algW* and Δ*prc* Δ*algW* strains (Fig. 2a). This could be due to the activation of an unknown protease in the absence of AlgW. Alternatively, the *algD* control region is the target of multiple regulators and perhaps one of these can still increase Φ(*algDp-lacZ*) expression slightly in Δ*algW* mutants (50–54).

**FIG 2.**
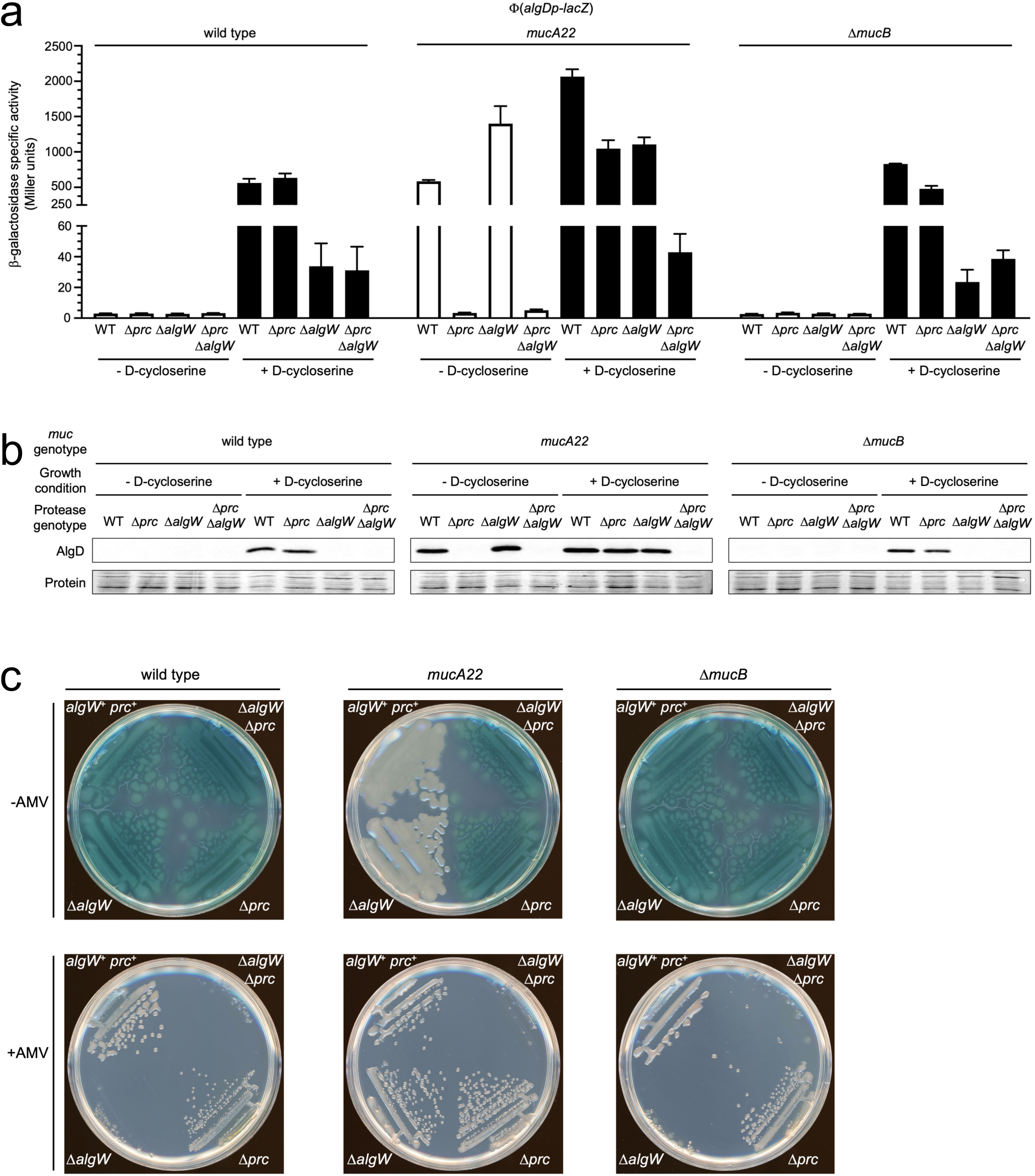
Comparison of the effects of Δ*prc* and Δ*algW* mutations on regulation of the *algD* operon and mucoidy. (a) Φ(*algDp-lacZ*) operon fusion expression. Strains with wild type *muc* genes (wild type), a *mucA22* mutation or a Δ*mucB* mutation were grown with or without exposure to D-cycloserine. Protease genotypes are shown below each bar (WT = wild type *prc*^+^ *algW*^+^). Error bars indicate the positive standard deviations from the means. (b) Anti-AlgD immunoblot analysis of one representative set of the triplicate cultures used to generate the data in panel a. (c) Phenotypes on Pseudomonas Isolation Agar with or without ammonium metavanadate (AMV). Immunoblots and agar plates are single representatives of several replicate experiments.

Next we analyzed *mucA22* strains. Constitutive high level Φ(*algDp-lacZ*) expression and AlgD protein production was completely suppressed in the Δ*prc* and Δ*prc* Δ*algW* strains, but not in the Δ*algW* strain (Fig. a,b). These data are consistent with Prc, but not AlgW, acting as the site-1 protease for MucA22 in the absence of inducing signals. Two previous reports concluded that an *algW* null mutation did suppress the constitutive *mucA22* phenotype in strain PAO1, which contrasts with our findings in strain PAK (39, 55). However, in the absence of inducing signals, AlgW is expected to be proteolytically inactive, which fits with our data showing that it does not contribute to the constitutive *mucA22* phenotype in PAK (16, 23). Also, our own analysis of PAO1 *mucA22* strains described below agreed with our findings in PAK.

After induction with D-cycloserine, individual Δ*algW* or Δ*prc* mutations only slightly reduced Φ(*algDp-lacZ*) expression in the *mucA22* strains, whereas it was almost completely abolished in a Δ*prc* Δ*algW* double mutant (Fig. 2a). Similarly, the AlgD protein level was unaffected by the individual mutations, but was undetectable in the double mutant (Fig. 2b). These data suggest that either Prc or AlgW is sufficient to act as the MucA22 site-1 protease in inducing conditions, which is consistent with both proteases being active in this situation.

We also examined the phenotypes of these strains on PIA +/- AMV. As expected, all of the *mucA^+^*strains were blue/green and non-mucoid in the absence of AMV (Fig. 2c). In the presence of AMV, the Δ*prc* mutation did not prevent the induction of a mucoid phenotype or loss of blue/green pigment as seen in the earlier experiment (Figs. 1c, 2c). In the original description of the PIA+AMV protocol, it was reported that a strain PAO1 Δ*algW* mutant was non mucoid on PIA+AMV (35). Our data in the PAK strain agreed with those findings. However, our PAK Δ*algW* and Δ*prc* Δ*algW* mutants were non-mucoid only because they did not grow on PIA+AMV (Fig. 2c). The failure of Δ*algW* mutants to grow was not noted in the original PAO1 study, so we were curious to know if it might be a PAK vs. PAO1 strain-specific difference. Therefore, we constructed Δ*prc* and Δ*algW* derivatives of strain MPAO1 and analyzed them on PIA +/- AMV. The results were indistinguishable from those in the PAK strain, with an MPAO1 Δ*algW* mutant failing to grow on PIA+AMV (Fig. S1). We suspected that Δ*algW* mutants cannot grow in these conditions because failure to induce the AlgU regulon results in the inability to mitigate what is now a lethal inducing stress. To test this further, we analyzed a PAK Δ*algU* mutant and found that it also failed to grow on PIA+AMV (Fig. S2).

In a *mucA22* background, the *ΔalgW* mutation did not suppress the constitutive mucoid phenotype in the absence of AMV, which is consistent with the Φ(*algDp-lacZ*) expression and AlgD protein data (Fig. 2). This was also corroborated by our analysis of *algW*^+^ and Δ*algW* MPAO1 strains with a *mucA22* mutation (Fig. S1). In the presence of AMV, the individual *Δprc* and *ΔalgW* mutations did not prevent the induction of mucoidy, whereas the Δ*prc* Δ*algW* double mutant did not grow (Fig. 2c). This is consistent with Prc or AlgW acting as the MucA22 site-1 protease in inducing conditions, when both proteases are active. It also further supports the suggestion that the inability to grow on PIA+AMV is due to the failure to induce the AlgU regulon.

All of the data described above suggest that in inducing conditions, where Prc and AlgW are both active, either is sufficient to act as the site 1 protease of MucA22. In contrast, in non-inducing conditions, where AlgW is inactive, Prc might be the only available site 1 protease for MucA22. However, as in all previous published studies, these genetic data alone are not sufficient to make the conclusion that Prc can degrade MucA22. This shortfall was addressed later with an *in vitro* proteolysis assay.

### A *mucB* null mutation does not induce mucoidy or make wild type MucA susceptible to Prc-dependent activation

It has been reported that *mucB* null mutations cause moderate constitutive activation of the AlgU regulon and mucoidy (21, 26, 28, 35, 37, 56–58). MucB protects the C-terminal domain of wild type MucA from proteolysis (23). Removal of MucB alone would not be expected to cause a mucoid phenotype because the AlgW site-1 protease would be inactive. However, it might cause a constitutive phenotype if removal of MucB from the C-terminal domain rendered wild type MucA susceptible to degradation by Prc. Therefore, we investigated the impact of Δ*algW* and Δ*prc* mutations in strains with a *mucB* in-frame deletion mutation. We found that the Δ*mucB* mutation did not cause constitutive induction of Φ(*algDp-lacZ*) expression, AlgD protein production, or mucoidy (Fig. 2). The constitutive phenotypes in the literature might be due to the use of *mucB* insertion mutations, or clinical isolates with *mucB* point mutations. These mutations could have polar effects on the downstream *mucC* and *mucD* genes, which can impact the AlgU regulon positively and negatively, respectively (59, 60). Polar effects are less likely with our *mucB* in-frame deletion mutation. In fact, the effects of Δ*algW* and Δ*prc* mutations in *mucA*^+^ *mucB*^+^ and *mucA*^+^ Δ*mucB* strains were indistinguishable, and so serve to further suggest that the periplasmic domain of wild type MucA is unlikely to be a substrate recognized by Prc, even when it is not protected from proteolysis by MucB (Fig. 2).

### Prc degrades the periplasmic domain of MucA22 *in vitro*

This and previous genetic studies have provided evidence that Prc might degrade MucA22, but the evidence is not unequivocal. Indirect effects of a *prc* null mutation could affect MucA22 stability, especially as MucA stability is known to be affected by cell envelope stress (37). Therefore, we wanted to test if Prc could degrade MucA or MucA22 *in vitro*. Attempted overproduction of Prc in the *E. coli* cytoplasm arrested growth immediately, so C-terminal His_6_ tagged Prc and an inactive Prc-S456A negative control (catalytic serine changed to alanine) were produced in the periplasm and purified using nickel agarose. The host strain had Δ*nlpI* and Δ*prc::kan* mutations to avoid contamination with *E. coli* Prc or its accessory protein NlpI (47). Overproduction of full length MucA and MucA22 was also lethal, so we purified maltose binding protein (MBP)-′MucA and -′MucA22 periplasmic domain fusion proteins (the periplasmic domain is the only part of MucA accessible to Prc in bacterial cells) The small sizes of the MucA and MucA22 periplasmic domains, approximately 90 and 40 amino acids, respectively, also meant that their purification was facilitated by fusion to the C-terminus of MBP.

Prc degraded MBP-′MucA22 whereas the Prc-S456A negative control did not (Fig. 3a). The fusion protein was not completely degraded, with the size of the reaction product suggesting that the MucA22 periplasmic domain might have been degraded while the MBP domain was left intact (Fig. 3a). Anti-MBP immunoblot analysis confirmed that the reaction product was a derivative of the MBP-′MucA22 protein. MBP-′MucA appeared to have been cleaved during production/purification, so that it was isolated as two major products corresponding in size to MBP-′MucA and MBP (Fig. 3b). We have observed a similar phenomenon with some other MBP fusion proteins in our laboratory (data not shown). Regardless, unlike MBP-′MucA22, the amount of full length MBP-‘MucA remained constant over time when it was incubated with Prc, suggesting that it was not a Prc substrate (Fig. 3b). The amount of the apparent MBP protein did increase slightly with Prc, but not with Prc-S456A. We suspect that this was because Prc had degraded minor intermediate proteins that ran between MBP-′MucA and MBP on the SDS-PAGE gels. Anti-MBP immunoblot analysis confirmed that these intermediate bands were derivatives of MBP-′MucA, and that they were eliminated upon incubation with Prc, but not with Prc-S456A (Fig. 3b). We hypothesize that these intermediate proteins are C-terminal truncation derivatives of MBP-′MucA, and that much like MBP-′MucA22, Prc degraded their MucA portions but left the MBP domain intact. This suggests that in addition to MucA22, other C-terminal truncations of MucA might be Prc substrates (see Discussion). Taken together, all of these *in vitro* data suggest that the periplasmic domain of MucA22 is a proteolytic substrate of Prc, whereas the full-length periplasmic domain of wild type MucA is not.

**FIG 3.**
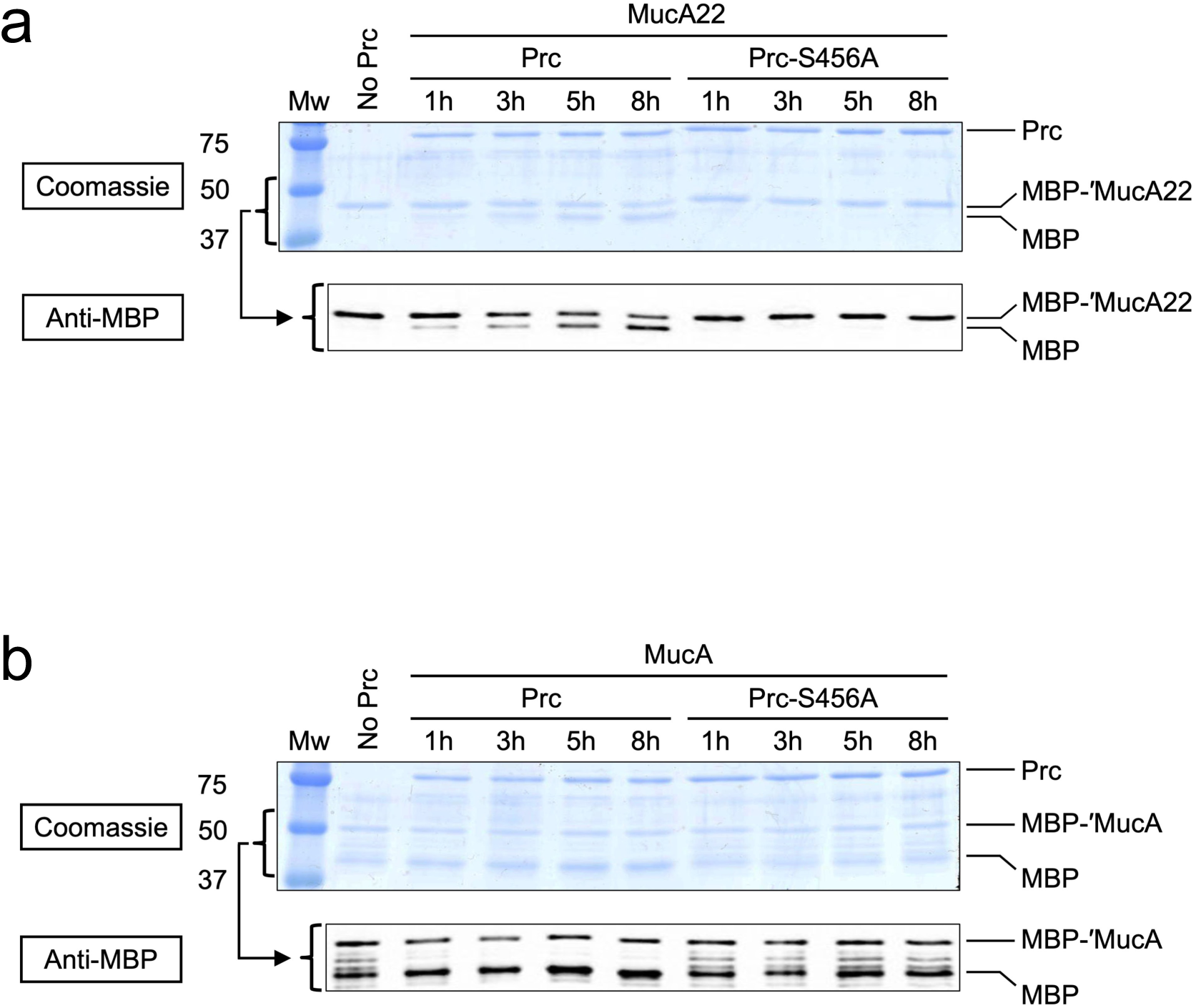
Analysis of MucA periplasmic domain cleavage *in vitro*. MBP-′MucA22 (a) or MBP-′MucA (b) proteins were incubated with Prc-His_6_ or Prc-S456-His_6_ proteins at 37°C for the indicated number of hours. For each panel the top image shows a Coomassie blue-stained SDS-PAGE gel with Mw markers sizes to the left (kDa), and the bottom image is anti-MBP immunoblot, with the bracket showing the corresponding regions of the two images. For wild type MucA, the MBP-′MucA protein purified as a mixture of mostly MBP-′MucA and MBP, along with some likely MBP-′MucA truncated intermediates visible as faint bands between MBP-′MucA and MBP on both the Coomassie blue-stained gel and anti-MBP immunoblot. Data presented are single representatives of several replicate experiments.

## DISCUSSION

Our laboratory has studied *P. aeruginosa* CTP family member CtpA, revealing that it plays a role strikingly similar to that played by Prc in *E. coli* (61–64)*. P. aeruginosa* has another CTP, which has been named Prc because it is a much closer homolog of *E. coli* Prc. This has motivated us to begin studying *P. aeruginosa* Prc, which has also been named AlgO because of its implication in controlling alginate biosynthesis, especially in clinical isolates with a mucoid phenotype caused by the *mucA22* mutation (32, 33). We found some of the literature unclear with regard to whether or not Prc might also play a role in strains with wild type MucA. We also noticed that some publications stated that Prc degrades MucA22, even though this had not been demonstrated. In fact, prior to this study, *P. aeruginosa* Prc had not been shown to degrade any protein. Therefore, we began our Prc studies with our own series of *in vivo* and *in vitro* experiments to investigate the impact of Prc on the regulation of alginate biosynthesis and mucoidy. Each one of our different experiments was in full agreement in support of the conclusions that AlgW is likely to be the only site-1 protease for wild type MucA in any condition, and that Prc is the only site-1 protease for MucA22 in normally non-inducing conditions. Notably, for the first time, we also showed that Prc is unable to degrade the periplasmic domain of wild type MucA, but that it does degrade the mutated periplasmic domain of MucA22 directly.

Only one of the two *P. aeruginosa* CTPs, Prc, had been investigated for its effect on the regulation of alginate biosynthesis. Therefore, we began by comparing the effects of both CTPs on activation of the AlgU regulon (Fig. 1). With wild type MucA, a *prc* null mutation did not reduce D-cycloserine-dependent induction of *algD* expression and AlgD protein production, or metavanadate-dependent induction of mucoidy. Three previous reports suggested that Prc might play a role with wild type MucA. One showed that a *prc* null mutation reduced D-cycloserine-dependent induction, but later quantitative analysis revealed that any effect was negligible (37, 38). Another concluded that Prc was essential for induction triggered by overproduction of the MucE protein, but an earlier report had reached the opposite conclusion (39, 40). A third reported that a strain PA14 *prc* transposon insertion mutant was non mucoid on PIA+AMV (36). However, the original PIA+AMV report showed that a strain PAO1 *prc* null mutant was still mucoid on PIA+AMV, in agreement with our findings. Perhaps the difference in PA14 was due to polar effects of the *prc* transposon insertion, or because the plates were incubated at 25°C, rather than 37°C as had been used in the original PAO1 study and our own experiments. Regardless, most reports agree with our findings that Prc does not significantly affect regulation in wild type MucA strains, a consensus that was also noted in an early review article (65). Like Δ*prc*, a Δ*ctpA* mutation also did not reduce induction in a strain with wild type MucA, showing that CtpA also plays no role in the wild type signal transduction pathway. This is consistent with the localization of CtpA in complex with its outer membrane lipoprotein partner LbcA, separating it from MucA in the inner membrane. However, we noticed that a Δ*ctpA* mutation did cause a small increase in D-cycloserine-dependent induction of *algD* expression and AlgD protein production. We suspect that this is because accumulation of the CtpA cell wall cross-link hydrolase substrates added to the cell wall stress caused by D-cycloserine (64).

Our genetic analysis corroborated conclusions from other studies that a *prc* null mutation suppresses the constitutive phenotype of a *mucA22* mutant (Figs. 2 and S1). There are two reports of an *algW* null mutation also suppressing the constitutive phenotype of a *mucA22* mutant (39, 55). However, that did not occur in our strains (Figs. 2 and S1). The suppression reported in the previous studies could not be complemented by *algW*^+^, perhaps suggesting that it had a different cause (39, 55). Our findings are consistent with the fact that AlgW is inactive in non-inducing conditions, because it must be engaged by mislocalized OMPs triggered by envelope stress in order to be active (16, 23). Therefore, removal of AlgW is not expected to impact the constitutive *mucA22* phenotype, unless the *mucA22* mutation itself triggers an AlgW-inducing signal. Our data suggests that it does not. In inducing conditions, removal of either Prc or AlgW alone did not abolish high level Φ(*algDp-lacZ*) expression, AlgD protein production, or mucoidy. This fits with AlgW having been activated in these conditions, so that both AlgW and Prc can act as the MucA22 site-1 protease. The AlgW cleavage site is still present in the truncated MucA22 protein (23). Therefore, only removal of both proteases abolishes activation. Taken together, all our data suggested that in contrast to wild type MucA, the MucA22 periplasmic domain might be a substrate for AlgW and Prc, but that Prc could be the only protease active to degrade it in normally-non-inducing conditions.

Our genetic analyses made two other findings of note. First, mutational removal of MucB, which shields the C-terminus of wild type MucA from proteolysis, did not trigger Prc-dependent activation (Fig. 2). This further supported the suggestion that wild type MucA cannot be cleaved by Prc. Some other studies had associated *mucB* null mutations with moderate increases of alginate production and mucoidy, but this might have been due to polar effects on the downstream *mucCD* genes triggering AlgW-activating conditions (21, 26, 28, 35, 37, 56–58). Consistent with that, the phenotype of one of those *mucB* mutants was not affected by a *prc* null mutation (32). Our second finding of note was that our data suggested that the inability to activate the AlgU regulon is lethal on PIA containing AMV, in both the PAK and PAO1 strains (Figs. 2 and S1). To our knowledge this has not been reported before, although previous reporting that an *algW* null mutant is non-mucoid on PIA+AMV is not incompatible with it being unable to grow (35). The most likely explanation is that one or more targets of AlgU are essential to mitigate the envelope stress caused by the combination of irgasan and AMV in these conditions. Strain PA14 *algW* and *algU* transposon insertion mutants did grow on PIA+AMV, but at a much lower temperature of 25°C that might have reduced the level of envelope stress (36).

MucA cleavage by AlgW has already been demonstrated *in vitro* (16, 23, 66). In contrast, cleavage by Prc had not been investigated prior to this study. Therefore, it is important that for the first time we showed that the periplasmic domain of MucA22 is a Prc substrate, whereas the periplasmic domain of MucA is not. This fits perfectly with the predictions made by all of our genetic analysis. There was also a tentative indication that Prc might degrade other C-terminal truncated derivatives of MucA that were isolated when we purified the MBP-′MucA protein (Fig. 3b). This idea is supported by the observation that in addition to *mucA22*, a *prc* null mutation suppressed the constitutive phenotypes caused by two other clinical *mucA* mutations that caused C-terminal truncations distinct from *mucA22* (32). This also raises the possibility that one role of Prc in *P. aeruginosa* might be to act as a clean-up protease that eliminates envelope proteins with aberrant C-termini. In fact, in *E. coli*, Prc degrades SsrA-tagged proteins in the periplasm (67, 68). SsrA-tags are co-translationally added to truncated C-termini caused by stalling of the translation machinery, and they mark proteins for destruction by several cytoplasmic proteases, or most likely by Prc in the cell envelope (69). Additional suggestion of a clean-up role for *E. coli* Prc comes from observations that it might prefer proteins with disordered C-termini (70). In *P. aeruginosa*, our ongoing investigations have not found any evidence that Prc works with an accessory outer membrane lipoprotein analogous to NlpI in *E. coli*, or LbcA for CtpA in *P. aeruginosa* (A. G. Sommerfield and A. J. Darwin, unpublished). Perhaps *E. coli* Prc can take on a clean-up role when in isolation, and an autolysin degradation role when in complex with NlpI. In contrast, these two roles might be divided between Prc and the LbcA-CtpA complex in *P. aeruginosa*.

In addition to AlgU-MucA, *P. aeruginosa* has several other sigma factor/anti-sigma factor regulatory systems, including cell-surface signaling systems involved in iron acquisition (71, 72). Prc has been implicated as a site-1 protease in some of these systems, in both *P. aeruginosa* and other *Pseudomonas* species (41–44). Prc has also been proposed to be a site-1 protease in the regulated intramembrane proteolysis of TcpP in *Vibrio cholerae* (73). However, Prc has not yet been shown to cleave any of those regulatory proteins directly. Even so, there is one example of Prc cleaving a transmembrane regulatory protein directly in *Xanthomonas campestris* pv. *campestris* (*74*). Therefore, while Prc is not involved in the wild type MucA signal transduction pathway, it remains possible that it has a direct role in activating other sigma factor/anti-sigma factor regulatory systems, in *P. aeruginosa* and beyond.

## MATERIAL AND METHODS

### Bacterial strains and standard growth conditions

*P. aeruginosa* strains and plasmids used in this study are listed in Table 1. *E. coli* K-12 strain SM10 was used for conjugation of plasmids into *P. aeruginosa* (75). For routine propagation, bacteria were grown in Luria-Bertani (LB) broth, composed of 1% (wt/vol) tryptone, 0.5% (wt/vol) yeast extract, 1% (wt/vol) NaCl, or on LB agar. To select for *P. aeruginosa* exconjugants after mating with *E. coli* donor strains, bacteria were recovered on Vogel-Bonner minimal agar with appropriate antibiotics (76).

**TABLE 1.**
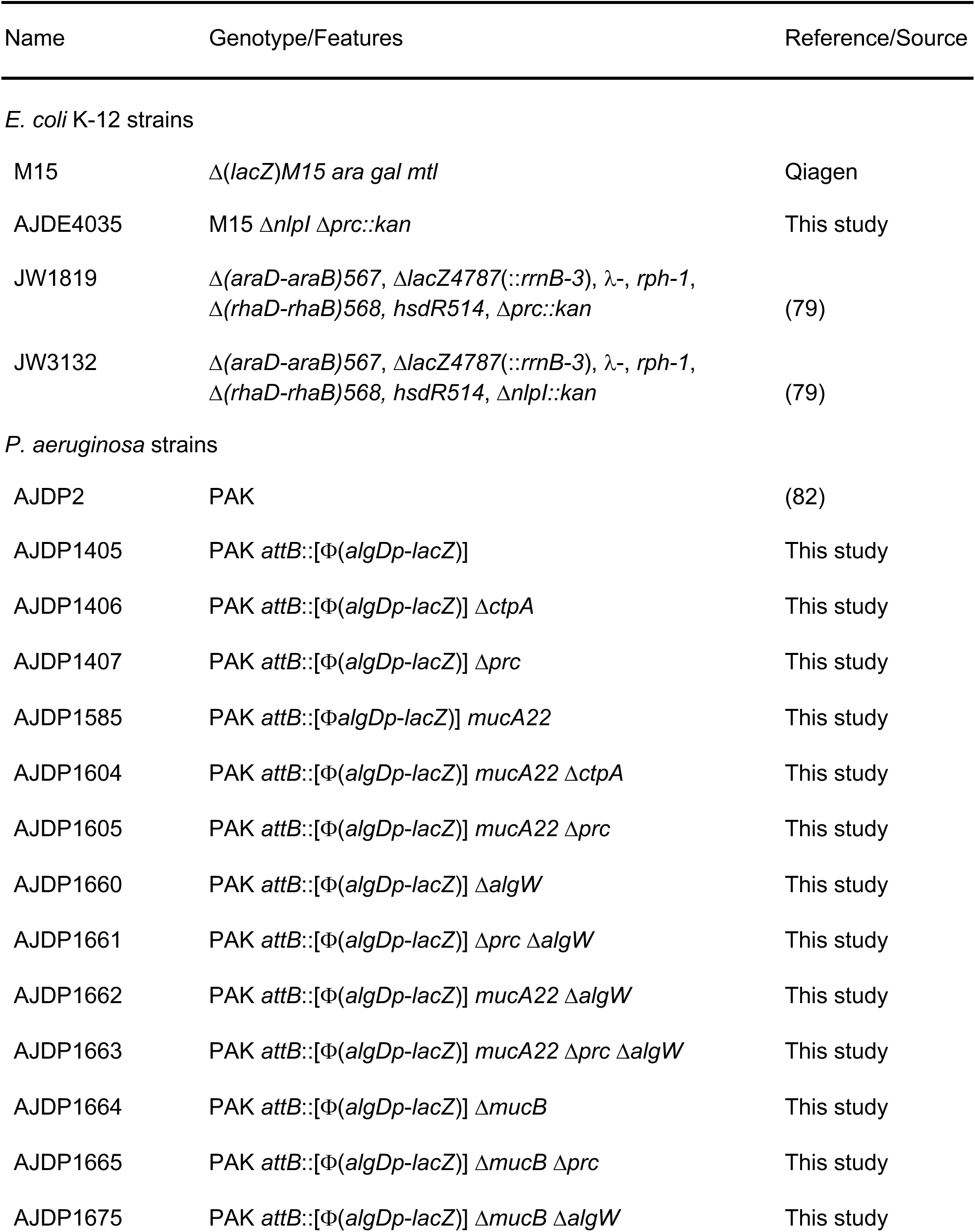
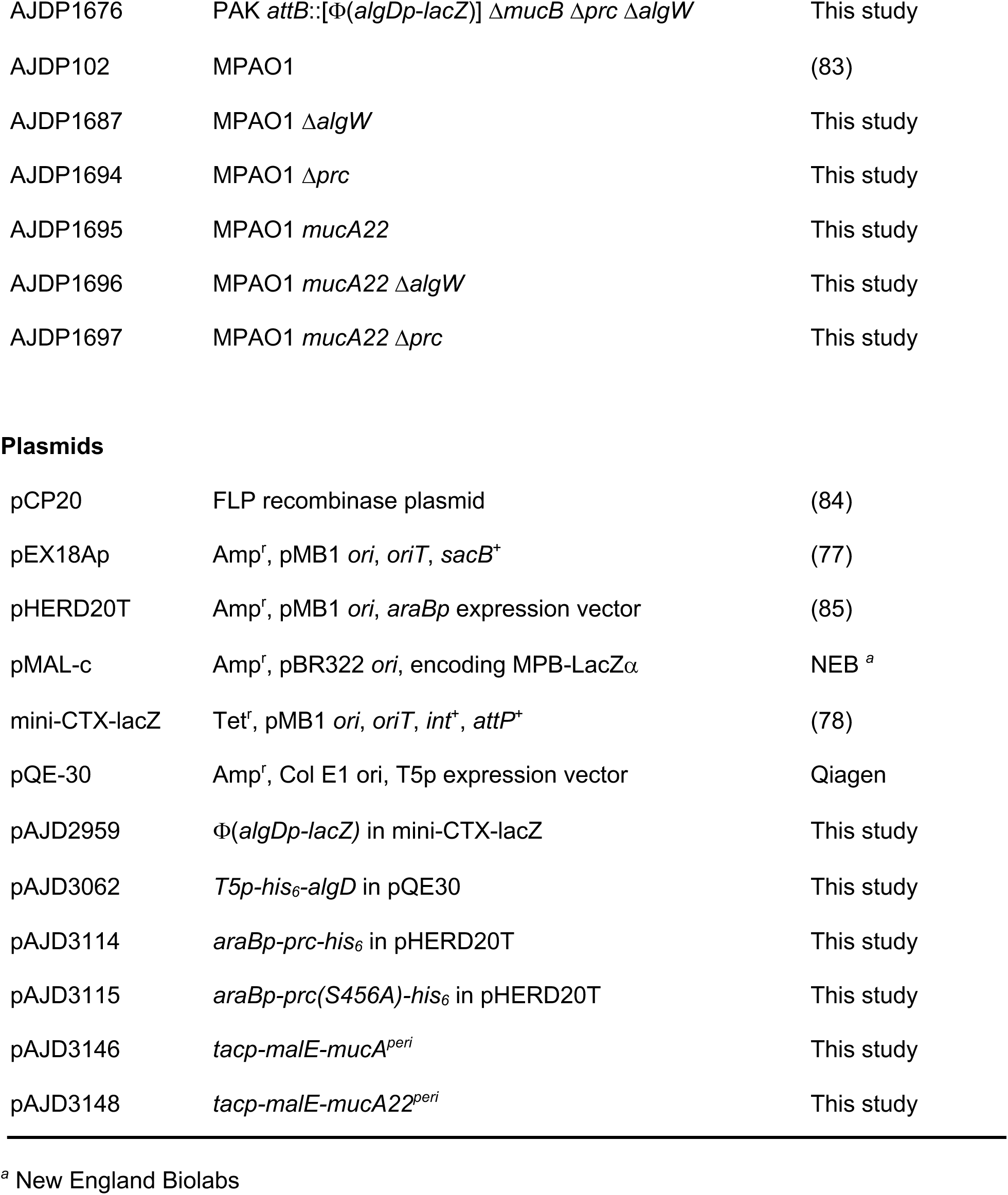
Strains and plasmids.

### Plasmid and strain construction

To construct strains with Δ*prc*, Δ*algW*, or Δ*mucB* in-frame deletion mutations, fragments of ∼0.55 kb corresponding to the regions upstream and downstream of the deletion site were amplified by PCR and cloned into pEX18Ap. These plasmids were integrated into the *P. aeruginosa* chromosome after conjugation from *E. coli* SM10 and sucrose-resistant carbenicillin-sensitive segregants were isolated on LB containing 10% sucrose without NaCl. Deletions were verified by PCR analysis of genomic DNA.

For the *mucA22* mutation, two ∼0.55-kb fragments flanking codon 144 of *mucA* were amplified by PCR. For each fragment, one of the primers incorporated a deletion of the first nucleotide of codon 144 to construct the *mucA22* frameshift. The fragments were joined in a PCR splicing by overlap extension (SOEing) reaction (50) via their overlapping regions around codon 144. The product was cloned into pEX18Ap and exchanged for the corresponding region of the *mucA* gene by integration, selection for sucrose-resistant segregants, and confirmation by colony PCR and DNA sequencing.

To construct single copy Φ(*algDp-lacZ)* fusion strains, the entire non-coding region upstream of *algD* was amplified by PCR and cloned into the mini-CTX-*lacZ* vector. This plasmid was inserted into the *attB* site after conjugation from *E. coli* SM10, and then vector backbone was then removed by FLP2-mediated excision (77). Strains were confirmed by PCR analysis of genomic DNA as described previously (78).

A derivative of *E. coli* strain M15 with *nlpI* and *prc* deletion mutations was constructed using mutations from the Keio collection (79). First the Δ*nlpI::kan* mutation was introduced into M15 by phage P1 *vir* transduction. The kanamycin cassette was removed by FLP2-mediated excision using plasmid pCP20 as described previously (11). The *nlpI* deletion was confirmed by colony PCR analysis. Next, the *Δprc::kan* mutation was introduced by phage P1 *vir* transduction and confirmed by colony PCR analysis.

A plasmid encoding His_6_-AlgD was constructed by amplifying the *algD* gene from PAK DNA, using a downstream primer that annealed immediately after the stop codon. This fragment was cloned into pQE-30 using restriction sites incorporated by the PCR primers.

To construct a *prc*-His_6_ expression plasmid, *prc* was amplified from strain PAK genomic DNA using an upstream primer that incorporated the ribosomal binding site from pQE30, and a downstream primer that included a region encoding the His_6_ tag, followed by a stop codon. To construct a similar plasmid encoding the catalytically inactive mutant Prc-S456A, two fragments flanking codon 456 of *prc* were amplified by PCR. For each fragment, one of the primers incorporated a mismatch at codon 456 to convert it to encode alanine. The fragments were joined in a PCR SOEing reaction via their overlapping regions around codon 456. The *prc*-His_6_ and *prc(S456A)*-His_6_ fragments were cloned into pHERD20T using restriction enzyme sites incorporated in the primers.

Plasmids encoding MBP-′MucA or -′MucA22 periplasmic domain fusion proteins were made by amplifying the *mucA* and *mucA22* periplasmic domains from genomic DNA. The upstream primer annealed immediately upstream of *mucA* codon 107 and incorporated an EcoRV site for in-frame fusion to the *malE* gene of plasmid pMAL-c. The downstream primers annealed immediately downstream of the *mucA* or *mucA22* stop codons and incorporated an XbaI site. The PCR fragments were digested with EcoRV/XbaI and ligated into plasmid pMAL-C plasmid that was digested with StuI/XbaI.

All plasmids constructed using PCR amplification were checked by DNA sequencing of the inserts to ensure that no spurious mutations had been introduced.

### Growth of bacteria +/- D-cycloserine

Bacteria were grown to saturation at 37°C in LB broth and diluted to an OD_600_ of 0.05 in 5 mL LB broth in 18mm test tubes. For each strain, duplicate cultures were grown at 37°C for 2 h in a roller drum, and then a final concentration of 200µg/mL of D-cycloserine was added to one of each duplicate. The cultures were grown for an additional 2 h before bacterial cells were collected by centrifugation for β-Galactosidase assays and immunoblot analysis.

### β-galactosidase activity assay

β-Galactosidase enzyme activity was determined at room temperature in permeabilized cells as described previously (80). Activities are expressed in arbitrary Miller units (81). Individual cultures were assayed in duplicate, and averages values from three independent cultures are reported.

### AlgD polyclonal antisera production and immunoblotting

*E. coli* strain M15 [pREP4] (Qiagen) containing plasmid pAJD3062 encoding His_6_-AlgD was grown in LB broth to mid-log phase at 37°C with aeration. Protein production was induced with 1 mM IPTG for 3 h at 37°C. The protein was purified under denaturing conditions by nickel-nitrilotriacetic acid (NTA)-agarose affinity chromatography as described by the manufacturer (Qiagen). A polyclonal rabbit antiserum was raised by Labcorp Early Development Laboratories Inc.

Samples were separated by SDS-PAGE and transferred to nitrocellulose by semi dry electroblotting. Chemiluminescent detection followed incubation with the AlgD polyclonal antiserum or anti-MBP monoclonal antibody (NEB), then goat anti-rabbit IgG (Sigma) or goat anti-mouse IgG (Sigma) horseradish peroxidase conjugates used at the manufacturers recommended dilution.

### Growth on *Pseudomonas* Isolation Agar (PIA) +/- ammonium metavanadate (AMV)

PIA containing 20 mL/L of glycerol was boiled before being autoclaved and cooled to 60°C. For inclusion of AMV, 35 mg of AMV was added to 14 mL of sterile water and boiled with a stir bar until the solution appeared light yellow. 12.5 ml of the 2.5 mg/mL AMV solution was added to 1 L of molten PIA for a final AMV concentration of 0.27 mM. PIA agar plates +/- AMV were dried in a laminar flow hood for 40 min and stored at 4°C for a maximum of one week. Saturated bacterial cultures were diluted to OD_600_ = 1 in LB broth and 5 µL of this suspension was streaked onto the agar plates, which were then incubated for 24 hours at 37°C.

### Protein purification and *in vitro* proteolysis assays

To purify Prc-His_6_ or Prc(S456A)-His_6_, *E. coli* strain AJDE4035 containing plasmid pAJD3114 or pAJD3115 was grown in 1L of LB broth at 37°C with aeration until the OD_600_ was between 0.6 and 1.0. Protein production was induced by adding 0.2% arabinose and incubating for 5 h at 37°C with aeration. Cells were resuspended in 50 mM NaH_2_PO_4_, 300 mM NaCl, 10 mM imidazole, pH 8.0 and incubated with 1 mg/mL lysozyme for 30 min on ice. Cells were disrupted by sonication and insoluble materials were removed by centrifugation. The proteins were purified under native conditions by NTA-agarose affinity chromatography, as recommended by the manufacturer (Qiagen). Elution was in 2 mL fractions using 50 mM NaH_2_PO4 and 300 mM NaCl buffer containing increasing concentrations of imidazole (50 to 250 mM, in 50 mM increments). For both proteins, fractions 3-10 were combined and used for the assays.

To purify MBP-¢MucA or MBP-¢MucA22 *E. coli* strain M15 containing plasmid pAJD3146 or pAJD3148 was grown in 1L of LB broth at 37°C with aeration until OD_600_ was between 0.6 and 1.0. Protein production was induced by adding 1 mM IPTG and incubating for 3 h at 37°C with aeration. Cells were resuspended in cold column buffer (20mM Tris-HCl, pH 7.4, 200mM NaCl, 1mM EDTA, pH 7.4). Roche complete protease inhibitor was added immediately as well as 1 mg/mL lysozyme. Cells were incubated on ice for 30 minutes, disrupted by sonication, and then insoluble material was removed by centrifugation. The supernatant was added to a column containing 2.5 mL amylose resin (NEB). The resin was washed with 50 mL of column buffer and proteins were eluted with 20 mL cold Column buffer containing 10mM maltose, in 1 mL fractions. For both proteins, fractions 2 and 3 were combined and used for the assays.

*In vitro* proteolysis reactions contained approximately 2 µM Prc-His_6_ or Prc-S456A-His_6_, and approximately 2 µM MBP-′MucA or MBP-′MucA22. Reactions were incubated at 37°C and aliquots were removed after 1, 3, 5, and 8 h and terminated by adding SDS-PAGE sample buffer and boiling. Proteins were separated by SDS-PAGE and stained with ProtoBlue Safe (National Diagnostics) or analyzed by anti-MBP immunoblot as described above.

## ACKNOWLEDGEMENTS

Research was supported by the National Institute of Allergy and Infectious Diseases (NIAID) of the National Institutes of Health (NIH), under award number R21AI151097 to A.J.D., and award number T32AI007180 also supported A.G.S. The content is solely the responsibility of the authors and does not necessarily represent the official views of the NIH.

---

**Supplemental Figure S1.**
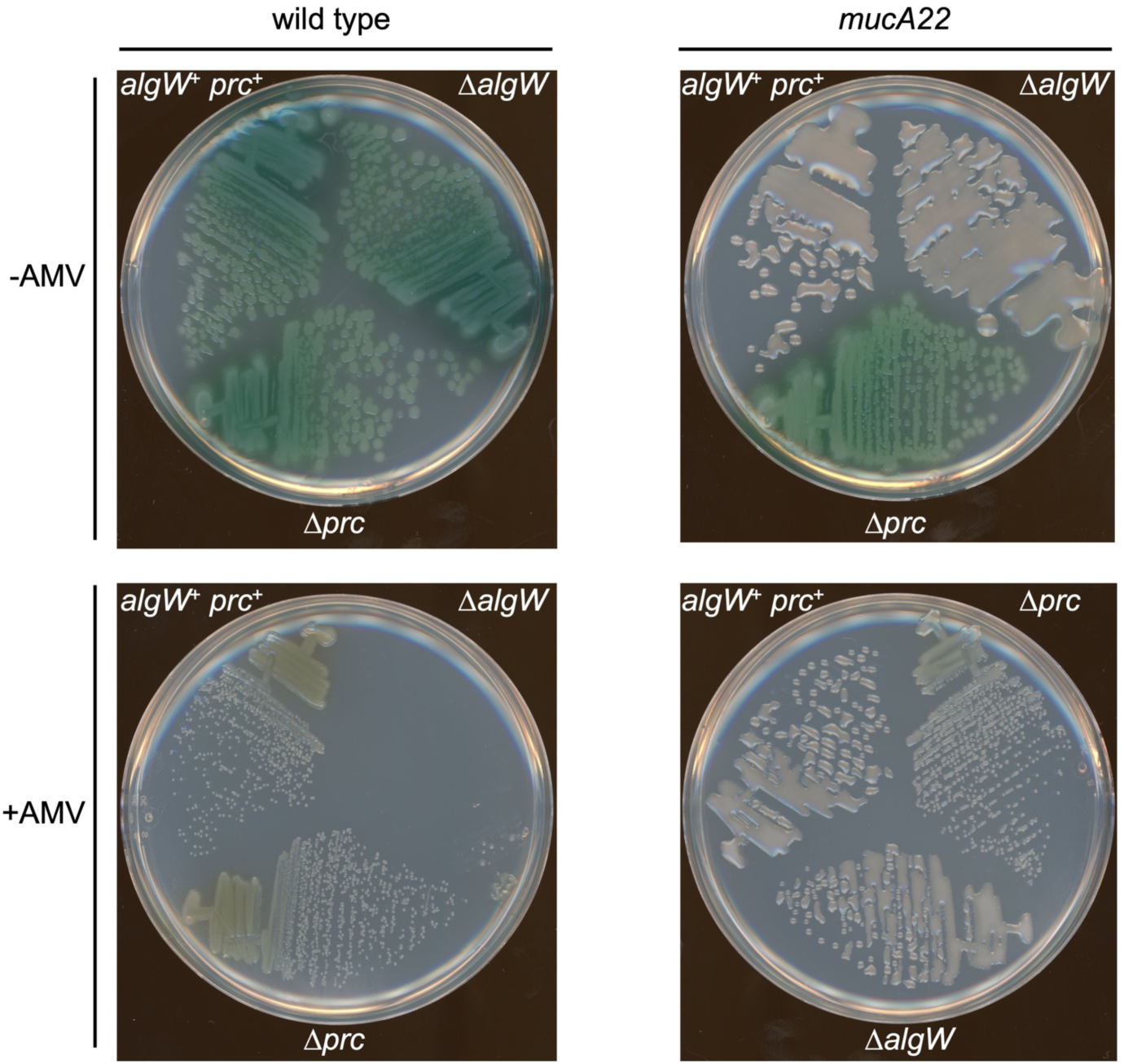
Analysis of strain MPAO1 derivative phenotypes on *Pseudomonas* Isolation Agar with or without ammonium metavanadate (AMV). Strains had the wild type *mucA* gene (wild type) or *mucA22*, as indicated at the top. Single representatives of several replicate experiments.

**Supplemental Figure S2.**
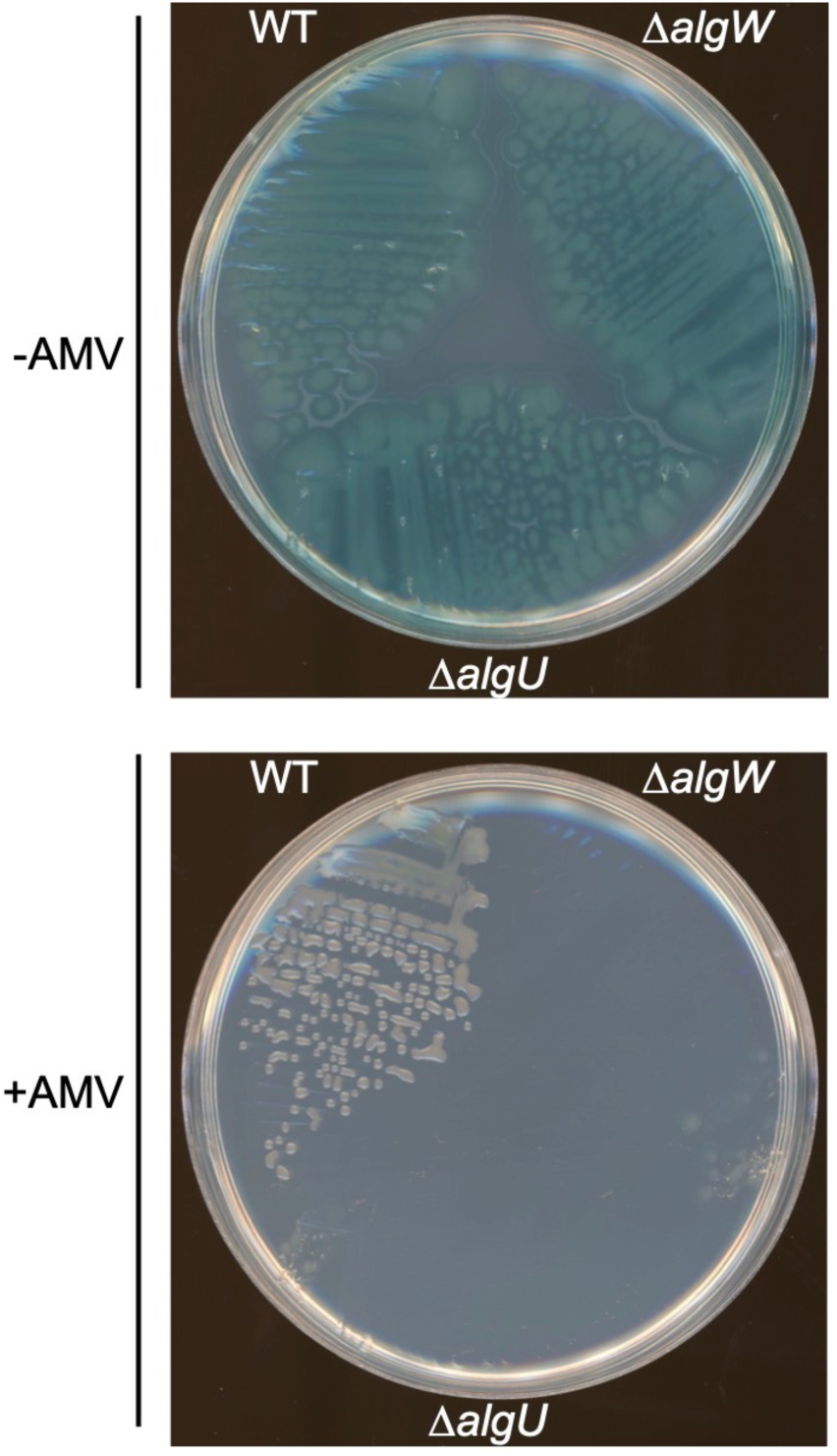
Comparison of PAK Δ*algW* and Δ*algU* phenotypes on Pseudomonas isolation agar with or without ammonium metavanadate (AMV). Strains had the wild type *mucA* gene (wild type) or *mucA22*, as indicated at the top. WT = *algW*^+^ *algU*^+^. Single representatives of several replicate experiments.

## REFERENCES

1. Moore NM, Flaws ML. 2011. Epidemiology and pathogenesis of *Pseudomonas aeruginosa* infections. Clin Lab Sci 24:43–46.

2. Kerem B, Rommens JM, Buchanan JA, Markiewicz D, Cox TK, Chakravarti A, Buchwald M, Tsui LC. 1989. Identification of the cystic fibrosis gene: genetic analysis. Science 245:1073–1080.

3. Rommens JM, Iannuzzi MC, Kerem B, Drumm ML, Melmer G, Dean M, Rozmahel R, Cole JL, Kennedy D, Hidaka N, et al. 1989. Identification of the cystic fibrosis gene: chromosome walking and jumping. Science 245:1059–1065.

4. Govan JR, Deretic V. 1996. Microbial pathogenesis in cystic fibrosis: mucoid *Pseudomonas aeruginosa* and *Burkholderia cepacia*. Microbiol Rev 60:539–574.

5. Henry RL, Mellis CM, Petrovic L. 1992. Mucoid *Pseudomonas aeruginosa* is a marker of poor survival in cystic fibrosis. Pediatr Pulmonol 12:158–161.

6. Leid JG, Willson CJ, Shirtliff ME, Hassett DJ, Parsek MR, Jeffers AK. 2005. The exopolysaccharide alginate protects *Pseudomonas aeruginosa* biofilm bacteria from IFN-gamma-mediated macrophage killing. J Immunol 175:7512–7518.

7. Govan JR, Fyfe JA. 1978. Mucoid *Pseudomonas aeruginosa* and cystic fibrosis: resistance of the mucoid from to carbenicillin, flucloxacillin and tobramycin and the isolation of mucoid variants in vitro. J Antimicrob Chemother 4:233–240.

8. Schwarzmann S, Boring JR. 1971. Antiphagocytic Effect of Slime from a Mucoid Strain of *Pseudomonas aeruginosa*. Infect Immun 3:762–767.

9. Wozniak DJ, Ohman DE. 1994. Transcriptional analysis of the *Pseudomonas aeruginosa* genes *algR*, *algB*, and *algD* reveals a hierarchy of alginate gene expression which is modulated by *algT*. J Bacteriol 176:6007–6014.

10. Chitnis CE, Ohman DE. 1993. Genetic analysis of the alginate biosynthetic gene cluster of *Pseudomonas aeruginosa* shows evidence of an operonic structure. Mol Microbiol 8:583–593.

11. Deretic V, Schurr MJ, Boucher JC, Martin DW. 1994. Conversion of *Pseudomonas aeruginosa* to mucoidy in cystic fibrosis: environmental stress and regulation of bacterial virulence by alternative sigma factors. J Bacteriol 176:2773–2780.

12. DeVries CA, Ohman DE. 1994. Mucoid-to-nonmucoid conversion in alginate-producing *Pseudomonas aeruginosa* often results from spontaneous mutations in *algT*, encoding a putative alternate sigma factor, and shows evidence for autoregulation. J Bacteriol 176:6677–6687.

13. Lonetto MA, Brown KL, Rudd KE, Buttner MJ. 1994. Analysis of the *Streptomyces coelicolor sigE* gene reveals the existence of a subfamily of eubacterial RNA polymerase sigma factors involved in the regulation of extracytoplasmic functions. Proc Natl Acad Sci U S A 91:7573–7577.

14. Hershberger CD, Ye RW, Parsek MR, Xie ZD, Chakrabarty AM. 1995. The *algT* (*algU*) gene of *Pseudomonas aeruginosa*, a key regulator involved in alginate biosynthesis, encodes an alternative sigma factor (sigma E). Proc Natl Acad Sci U S A 92:7941–7945.

15. Ades SE. 2004. Control of the alternative sigma factor sigmaE in *Escherichia coli*. Curr Opin Microbiol 7:157–162.

16. Lima S, Guo MS, Chaba R, Gross CA, Sauer RT. 2013. Dual molecular signals mediate the bacterial response to outer-membrane stress. Science 340:837–841.

17. Alba BM, Leeds JA, Onufryk C, Lu CZ, Gross CA. 2002. DegS and YaeL participate sequentially in the cleavage of RseA to activate the sigma(E)-dependent extracytoplasmic stress response. Genes Dev 16:2156–2168.

18. Alba BM, Zhong HJ, Pelayo JC, Gross CA. 2001. *degS* (*hhoB*) is an essential *Escherichia coli* gene whose indispensable function is to provide sigma (E) activity. Mol Microbiol 40:1323–1333.

19. Kanehara K, Ito K, Akiyama Y. 2002. YaeL (EcfE) activates the sigma(E) pathway of stress response through a site-2 cleavage of anti-sigma(E), RseA. Genes Dev 16:2147–2155.

20. Walsh NP, Alba BM, Bose B, Gross CA, Sauer RT. 2003. OMP peptide signals initiate the envelope-stress response by activating DegS protease via relief of inhibition mediated by its PDZ domain. Cell 113:61–71.

21. Schurr MJ, Yu H, Martinez-Salazar JM, Boucher JC, Deretic V. 1996. Control of AlgU, a member of the sigma E-like family of stress sigma factors, by the negative regulators MucA and MucB and *Pseudomonas aeruginosa* conversion to mucoidy in cystic fibrosis. J Bacteriol 178:4997–5004.

22. Rowen DW, Deretic V. 2000. Membrane-to-cytosol redistribution of ECF sigma factor AlgU and conversion to mucoidy in *Pseudomonas aeruginosa* isolates from cystic fibrosis patients. Mol Microbiol 36:314–327.

23. Cezairliyan BO, Sauer RT. 2009. Control of *Pseudomonas aeruginosa* AlgW protease cleavage of MucA by peptide signals and MucB. Mol Microbiol 72:368–379.

24. Martin DW, Schurr MJ, Mudd MH, Govan JR, Holloway BW, Deretic V. 1993. Mechanism of conversion to mucoidy in *Pseudomonas aeruginosa* infecting cystic fibrosis patients. Proc Natl Acad Sci U S A 90:8377–8381.

25. Boucher JC, Yu H, Mudd MH, Deretic V. 1997. Mucoid *Pseudomonas aeruginosa* in cystic fibrosis: characterization of *muc* mutations in clinical isolates and analysis of clearance in a mouse model of respiratory infection. Infect Immun 65:3838–3846.

26. Ciofu O, Lee B, Johannesson M, Hermansen NO, Meyer P, Hoiby N. 2008. Investigation of the *algT* operon sequence in mucoid and non-mucoid *Pseudomonas aeruginosa* isolates from 115 Scandinavian patients with cystic fibrosis and in 88 *in vitro* non-mucoid revertants. Microbiology (Reading) 154:103–113.

27. Pulcrano G, Iula DV, Raia V, Rossano F, Catania MR. 2012. Different mutations in *mucA* gene of *Pseudomonas aeruginosa* mucoid strains in cystic fibrosis patients and their effect on *algU* gene expression. New Microbiol 35:295–305.

28. Candido Cacador N, Paulino da Costa Capizzani C, Gomes Monteiro Marin Torres LA, Galetti R, Ciofu O, da Costa Darini AL, Hoiby N. 2018. Adaptation of P*seudomonas aeruginosa* to the chronic phenotype by mutations in the *algTmucABD* operon in isolates from Brazilian cystic fibrosis patients. PLoS One 13:e0208013.

29. Hwang W, Yong JH, Min KB, Lee KM, Pascoe B, Sheppard SK, Yoon SS. 2021. Genome-wide association study of signature genetic alterations among *Pseudomonas aeruginosa* cystic fibrosis isolates. PLoS Pathog 17:e1009681.

30. Anthony M, Rose B, Pegler MB, Elkins M, Service H, Thamotharampillai K, Watson J, Robinson M, Bye P, Merlino J, Harbour C. 2002. Genetic analysis of *Pseudomonas aeruginosa* isolates from the sputa of Australian adult cystic fibrosis patients. J Clin Microbiol 40:2772–2778.

31. Ryall B, Carrara M, Zlosnik JE, Behrends V, Lee X, Wong Z, Lougheed KE, Williams HD. 2014. The mucoid switch in *Pseudomonas aeruginosa* represses quorum sensing systems and leads to complex changes to stationary phase virulence factor regulation. PLoS One 9:e96166.

32. Reiling SA, Jansen JA, Henley BJ, Singh S, Chattin C, Chandler M, Rowen DW. 2005. Prc protease promotes mucoidy in *mucA* mutants of *Pseudomonas aeruginosa*. Microbiology (Reading) 151:2251–2261.

33. Sautter R, Ramos D, Schneper L, Ciofu O, Wassermann T, Koh CL, Heydorn A, Hentzer M, Hoiby N, Kharazmi A, Molin S, Devries CA, Ohman DE, Mathee K. 2012. A complex multilevel attack on *Pseudomonas aeruginosa algT/U* expression and *algT/U* activity results in the loss of alginate production. Gene 498:242–253.

34. Sommerfield AG, Darwin AJ. 2022. Bacterial Carboxyl-Terminal Processing Proteases Play Critical Roles in the Cell Envelope and Beyond. J Bacteriol 204:e0062821.

35. Damron FH, Davis MR, Jr., Withers TR, Ernst RK, Goldberg JB, Yu G, Yu HD. 2011. Vanadate and triclosan synergistically induce alginate production by *Pseudomonas aeruginosa* strain PAO1. Mol Microbiol 81:554–570.

36. Damron FH, Barbier M, McKenney ES, Schurr MJ, Goldberg JB. 2013. Genes required for and effects of alginate overproduction induced by growth of *Pseudomonas aeruginosa* on *Pseudomonas* isolation agar supplemented with ammonium metavanadate. J Bacteriol 195:4020–4036.

37. Wood LF, Leech AJ, Ohman DE. 2006. Cell wall-inhibitory antibiotics activate the alginate biosynthesis operon in *Pseudomonas aeruginosa*: Roles of sigma (AlgT) and the AlgW and Prc proteases. Mol Microbiol 62:412–426.

38. Wood LF, Ohman DE. 2009. Use of cell wall stress to characterize sigma 22 (AlgT/U) activation by regulated proteolysis and its regulon in *Pseudomonas aeruginosa*. Mol Microbiol 72:183–201.

39. Delgado C, Florez L, Lollett I, Lopez C, Kangeyan S, Kumari H, Stylianou M, Smiddy RJ, Schneper L, Sautter RT, Smith D, Szatmari G, Mathee K. 2018. *Pseudomonas aeruginosa* Regulated Intramembrane Proteolysis: Protease MucP Can Overcome Mutations in the AlgO Periplasmic Protease To Restore Alginate Production in Nonmucoid Revertants. J Bacteriol 200:e00215–00218.

40. Qiu D, Eisinger VM, Rowen DW, Yu HD. 2007. Regulated proteolysis controls mucoid conversion in *Pseudomonas aeruginosa*. Proc Natl Acad Sci U S A 104:8107–8112.

41. Bastiaansen KC, Civantos C, Bitter W, Llamas MA. 2017. New Insights into the Regulation of Cell-Surface Signaling Activity Acquired from a Mutagenesis Screen of the *Pseudomonas putida* IutY Sigma/Anti-Sigma Factor. Front Microbiol 8:747.

42. Bastiaansen KC, Ibanez A, Ramos JL, Bitter W, Llamas MA. 2014. The Prc and RseP proteases control bacterial cell-surface signalling activity. Environ Microbiol 16:2433–2443.

43. Otero-Asman JR, Garcia-Garcia AI, Civantos C, Quesada JM, Llamas MA. 2019. *Pseudomonas aeruginosa* possesses three distinct systems for sensing and using the host molecule haem. Environ Microbiol 21:4629–4647.

44. Otero-Asman JR, Sanchez-Jimenez A, Bastiaansen KC, Wettstadt S, Civantos C, Garcia-Puente A, Bitter W, Llamas MA. 2023. The Prc and CtpA proteases modulate cell-surface signaling activity and virulence in *Pseudomonas aeruginosa*. iScience 26:107216.

45. Jeon WJ, Cho H. 2022. A Cell Wall Hydrolase MepH Is Negatively Regulated by Proteolysis Involving Prc and NlpI in *Escherichia coli*. Front Microbiol 13:878049.

46. Kim YJ, Choi BJ, Park SH, Lee HB, Son JE, Choi U, Chi WJ, Lee CR. 2021. Distinct Amino Acid Availability-Dependent Regulatory Mechanisms of MepS and MepM Levels in *Escherichia coli*. Front Microbiol 12:677739.

47. Singh SK, Parveen S, SaiSree L, Reddy M. 2015. Regulated proteolysis of a cross-link-specific peptidoglycan hydrolase contributes to bacterial morphogenesis. Proc Natl Acad Sci U S A 112:10956–10961.

48. Hsu PC, Chen CS, Wang S, Hashimoto M, Huang WC, Teng CH. 2020. Identification of MltG as a Prc Protease Substrate Whose Dysregulation Contributes to the Conditional Growth Defect of Prc-Deficient *Escherichia coli*. Front Microbiol 11:2000.

49. Yakhnina AA, Bernhardt TG. 2020. The Tol-Pal system is required for peptidoglycan-cleaving enzymes to complete bacterial cell division. Proc Natl Acad Sci U S A 117:6777–6783.

50. Deretic V, Konyecsni WM. 1990. A procaryotic regulatory factor with a histone H1-like carboxy-terminal domain: clonal variation of repeats within *algP*, a gene involved in regulation of mucoidy in *Pseudomonas aeruginosa*. J Bacteriol 172:5544–5554.

51. Kato J, Misra TK, Chakrabarty AM. 1990. AlgR3, a protein resembling eukaryotic histone H1, regulates alginate synthesis in *Pseudomonas aeruginosa*. Proc Natl Acad Sci U S A 87:2887–2891.

52. Konyecsni WM, Deretic V. 1990. DNA sequence and expression analysis of *algP* and *algQ*, components of the multigene system transcriptionally regulating mucoidy in *Pseudomonas aeruginosa*: *algP* contains multiple direct repeats. J Bacteriol 172:2511–2520.

53. Mohr CD, Hibler NS, Deretic V. 1991. AlgR, a response regulator controlling mucoidy in *Pseudomonas aeruginosa*, binds to the FUS sites of the *algD* promoter located unusually far upstream from the mRNA start site. J Bacteriol 173:5136–5143.

54. Xu B, Soukup RJ, Jones CJ, Fishel R, Wozniak DJ. 2016. *Pseudomonas aeruginosa* AmrZ Binds to Four Sites in the *algD* Promoter, Inducing DNA-AmrZ Complex Formation and Transcriptional Activation. J Bacteriol 198:2673–2681.

55. Pandey S, Delgado C, Kumari H, Florez L, Mathee K. 2018. Outer-membrane protein LptD (PA0595) plays a role in the regulation of alginate synthesis in *Pseudomonas aeruginosa*. J Med Microbiol 67:1139–1156.

56. Wiens JR, Vasil AI, Schurr MJ, Vasil ML. 2014. Iron-regulated expression of alginate production, mucoid phenotype, and biofilm formation by *Pseudomonas aeruginosa*. mBio 5:e01010–01013.

57. Schweizer HP, Po C, Bacic MK. 1995. Identification of *Pseudomonas aeruginosa glpM*, whose gene product is required for efficient alginate biosynthesis from various carbon sources. J Bacteriol 177:4801–4804.

58. Martin DW, Schurr MJ, Mudd MH, Deretic V. 1993. Differentiation of *Pseudomonas aeruginosa* into the alginate-producing form: inactivation of mucB causes conversion to mucoidy. Mol Microbiol 9:497–506.

59. Boucher JC, Schurr MJ, Yu H, Rowen DW, Deretic V. 1997. *Pseudomonas aeruginosa* in cystic fibrosis: role of *mucC* in the regulation of alginate production and stress sensitivity. Microbiology (Reading) 143:3473–3480.

60. Damron FH, Yu HD. 2011. *Pseudomonas aeruginosa* MucD regulates the alginate pathway through activation of MucA degradation via MucP proteolytic activity. J Bacteriol 193:286–291.

61. Chakraborty D, Darwin AJ. 2021. Direct and indirect interactions promote complexes of the lipoprotein LbcA, the CtpA protease and its substrates, and other cell wall proteins in *Pseudomonas aeruginosa*. J Bacteriol:e0039321.

62. Chung S, Darwin AJ. 2020. The C-terminus of substrates is critical but not sufficient for their degradation by the *Pseudomonas aeruginosa* CtpA protease. J Bacteriol:e0017420.

63. Seo J, Darwin AJ. 2013. The *Pseudomonas aeruginosa* periplasmic protease CtpA can affect systems that impact its ability to mount both acute and chronic infections. Infect Immun 81:4561–4570.

64. Srivastava D, Seo J, Rimal B, Kim SJ, Zhen S, Darwin AJ. 2018. A Proteolytic Complex Targets Multiple Cell Wall Hydrolases in *Pseudomonas aeruginosa*. mBio 9:e00972–00918.

65. Damron FH, Goldberg JB. 2012. Proteolytic regulation of alginate overproduction in *Pseudomonas aeruginosa*. Mol Microbiol 84:595–607.

66. Li T, He L, Li C, Kang M, Song Y, Zhu Y, Shen Y, Zhao N, Zhao C, Yang J, Huang Q, Mou X, Tong A, Yang J, Wang Z, Ji C, Li H, Tang H, Bao R. 2020. Molecular basis of the lipid-induced MucA-MucB dissociation in *Pseudomonas aeruginosa*. Commun Biol 3:418.

67. Spiers A, Lamb HK, Cocklin S, Wheeler KA, Budworth J, Dodds AL, Pallen MJ, Maskell DJ, Charles IG, Hawkins AR. 2002. PDZ domains facilitate binding of high temperature requirement protease A (HtrA) and tail-specific protease (Tsp) to heterologous substrates through recognition of the small stable RNA A (*ssrA*)-encoded peptide. J Biol Chem 277:39443–39449.

68. Keiler KC, Waller PR, Sauer RT. 1996. Role of a peptide tagging system in degradation of proteins synthesized from damaged messenger RNA. Science 271:990–993.

69. Fritze J, Zhang M, Luo Q, Lu X. 2020. An overview of the bacterial SsrA system modulating intracellular protein levels and activities. Appl Microbiol Biotechnol 104:5229–5241.

70. Keiler KC, Silber KR, Downard KM, Papayannopoulos IA, Biemann K, Sauer RT. 1995. C-terminal specific protein degradation: activity and substrate specificity of the Tsp protease. Protein Sci 4:1507–1515.

71. Chevalier S, Bouffartigues E, Bazire A, Tahrioui A, Duchesne R, Tortuel D, Maillot O, Clamens T, Orange N, Feuilloley MGJ, Lesouhaitier O, Dufour A, Cornelis P. 2019. Extracytoplasmic function sigma factors in *Pseudomonas aeruginosa*. Biochim Biophys Acta Gene Regul Mech 1862:706–721.

72. Llamas MA, Mooij MJ, Sparrius M, Vandenbroucke-Grauls CM, Ratledge C, Bitter W. 2008. Characterization of five novel *Pseudomonas aeruginosa* cell-surface signalling systems. Mol Microbiol 67:458–472.

73. Teoh WP, Matson JS, DiRita VJ. 2015. Regulated intramembrane proteolysis of the virulence activator TcpP in *Vibrio cholerae* is initiated by the tail-specific protease (Tsp). Mol Microbiol 97:822–831.

74. Deng CY, Zhang H, Wu Y, Ding LL, Pan Y, Sun ST, Li YJ, Wang L, Qian W. 2018. Proteolysis of histidine kinase VgrS inhibits its autophosphorylation and promotes osmostress resistance in *Xanthomonas campestris*. Nat Commun 9:4791.

75. Miller VL, Mekalanos JJ. 1988. A novel suicide vector and its use in construction of insertion mutations: osmoregulation of outer membrane proteins and virulence determinants in *Vibrio cholerae* requires *toxR*. J Bacteriol 170:2575–2583.

76. Vogel HJ, Bonner DM. 1956. Acetylornithinase of *Escherichia coli*: partial purification and some properties. J Biol Chem 218:97–106.

77. Hoang TT, Karkhoff-Schweizer RR, Kutchma AJ, Schweizer HP. 1998. A broad-host-range Flp-FRT recombination system for site-specific excision of chromosomally-located DNA sequences: application for isolation of unmarked *Pseudomonas aeruginosa* mutants. Gene 212:77–86.

78. Becher A, Schweizer HP. 2000. Integration-proficient *Pseudomonas aeruginosa* vectors for isolation of single-copy chromosomal *lacZ* and *lux* gene fusions. Biotechniques 29:948–950, 952.

79. Baba T, Ara T, Hasegawa M, Takai Y, Okumura Y, Baba M, Datsenko KA, Tomita M, Wanner BL, Mori H. 2006. Construction of *Escherichia coli* K-12 in-frame, single-gene knockout mutants: the Keio collection. Mol Syst Biol 2:2006 0008.

80. Maloy SR, Stewart VJ, Taylor RK. 1996. Genetic Analysis of Pathogenic Bacteria: A Laboratory Manual. Cold Spring Harbor Laboratory Press, Plainview, NY.

81. Miller JH. 1972. Experiments in Molecular Genetics. Cold Spring Harbor Laboratory, Cold Spring Harbor, New York.

82. Strom MS, Lory S. 1986. Cloning and expression of the pilin gene of *Pseudomonas aeruginosa PAK* in Escherichia coli. J Bacteriol 165:367–372.

83. Jacobs MA, Alwood A, Thaipisuttikul I, Spencer D, Haugen E, Ernst S, Will O, Kaul R, Raymond C, Levy R, Chun-Rong L, Guenthner D, Bovee D, Olson MV, Manoil C. 2003. Comprehensive transposon mutant library of *Pseudomonas aeruginosa*. Proc Natl Acad Sci U S A 100:14339–14344.

84. Cherepanov PP, Wackernagel W. 1995. Gene disruption in *Escherichia coli*: TcR and KmR cassettes with the option of Flp-catalyzed excision of the antibiotic-resistance determinant. Gene 158:9–14.

85. Qiu D, Damron FH, Mima T, Schweizer HP, Yu HD. 2008. PBAD-based shuttle vectors for functional analysis of toxic and highly regulated genes in *Pseudomonas* and *Burkholderia* spp. and other bacteria. Appl Environ Microbiol 74:7422–7426.

